# Cryo-EM Structures of the N501Y SARS-CoV-2 Spike Protein in Complex with ACE2 and Two Potent Neutralizing Antibodies

**DOI:** 10.1101/2021.01.11.426269

**Authors:** Xing Zhu, Dhiraj Mannar, Shanti S. Srivastava, Alison M. Berezuk, Jean-Philippe Demers, James W Saville, Karoline Leopold, Wei Li, Dimiter S. Dimitrov, Katharine S. Tuttle, Steven Zhou, Sagar Chittori, Sriram Subramaniam

## Abstract

The recently reported “UK variant” of SARS-CoV-2 is thought to be more infectious than previously circulating strains as a result of several changes, including the N501Y mutation. We present a 2.9-Å resolution cryo-EM structure of the complex between the ACE2 receptor and N501Y spike protein ectodomains that shows Y501 inserted into a cavity at the binding interface near Y41 of ACE2. The additional interactions result in increased affinity of ACE2 for the N501Y mutant, accounting for its increased infectivity. However, this mutation does not result in large structural changes, enabling important neutralization epitopes to be retained in the spike receptor binding domain. We confirmed this through biophysical assays and by determining cryo-EM structures of spike protein ectodomains bound to two representative potent neutralizing antibody fragments.

**Short summary:** The N501Y mutation found in the coronavirus UK variant increases infectivity but some neutralizing antibodies can still bind.

## Introduction

The rapid international spread of severe acute respiratory syndrome coronavirus 2 (SARS-CoV-2), the causative agent of COVID-19, is associated with numerous mutations that alter viral fitness. Mutations have been documented in all four structural proteins encoded by the viral genome including the small envelope glycoprotein (E), membrane glycoprotein (M), nucleocapsid (N) protein, and the spike (S) protein. The most prominent mutations are in the spike protein, which mediates entry of the virus into cells by engaging with the angiotensin-converting enzyme-2 (ACE2) receptor. Several structures of SARS-CoV-2 spike protein variants in pre- and postfusion conformations have been reported, including complexes with ACE2 and a variety of antibodies (*1–13*). Mutations that emerge in the receptor binding domain (RBD) of the spike protein are especially of interest given their high potential to alter the kinetics and strength of interaction of the virus with target cells. These mutations could also affect the binding of antibodies capable of binding and blocking engagement of the virus with ACE2.

In December 2020, new variants of SARS-CoV-2 carrying several mutations in the spike protein were documented in the UK (SARS-CoV-2 VOC202012/01) and South Africa (501Y.V2) (*14, 15*). Early epidemiological and clinical findings have indicated that these variants show increased transmissibility in the population (*16*). Despite being phylogenetically distinct, a common feature of both UK and South African variants is the mutation of residue 501 in the RBD from Asn to Tyr (N501Y). X-ray crystallography and cryo-electron microscopy (cryo-EM) structural studies have identified N501 as a key residue in the spike protein at the interface between RBD and ACE2 that is involved in critical contacts with several ACE2 residues (*5, 6, 10, 13*). Studies carried out in a mouse model before the identification of the new UK variant have suggested that mutations of residue 501 could be linked to increased receptor binding and infectivity (*17, 18*). Understanding the impact of N501Y on antibody neutralization, ACE2 binding, and viral entry is therefore of fundamental interest in the efforts to prevent the spread of COVID-19.

## Results

### Visualization of Y501 in contact with ACE2

To understand the structural effects of the N501Y mutation on ACE2 binding, we expressed and purified spike (S) protein ectodomains with and without the N501Y mutation in Expi293 cells, and conducted microscopy studies on the ACE2–spike complexes. A cryo-EM structure of the spike protein ectodomain with the N501Y mutation was obtained at an average resolution of ~2.8 Å (Figure S1). The structure shows no significant global changes in secondary or quaternary structure as a result of the mutation when compared to the previously published structure of the spike protein ectodomain with an Asn residue at position 501 (referred to here as the “unmutated” form; Figure S2) (*7*).

Cryo-EM structural analysis of the complex formed between the N501Y spike protein ectodomain and the ACE2 receptor ectodomain provides a detailed glimpse of both the overall structure of the receptor and the binding interface between the RBD and ACE2 (Figures 1, S3). The ACE2 receptor is bound to the “up” position of the RBD (Figure 1A). The overall structure of the complex was determined at a global resolution of 2.9 Å. Local refinement of the RBD-ACE2 interface improves the local resolution at the binding interface to ~3.3 Å (Figure 1B), resulting in unambiguous delineation of the Y501 side chain and other residues in the vicinity (Figure 1C). The overall structure at the binding site is almost identical to that of the unmutated version (Figure 1D) (*7*), with the exception of local rearrangements that result in the aromatic ring of Y501 being accommodated in a cavity that is sandwiched between Y41 and K353 from the ACE2 receptor (Figure 1E). Y501 in the spike protein and Y41 in the ACE2 receptor form a perpendicular y-shaped π–π stacking interaction (*19*).

**Fig. 1.**
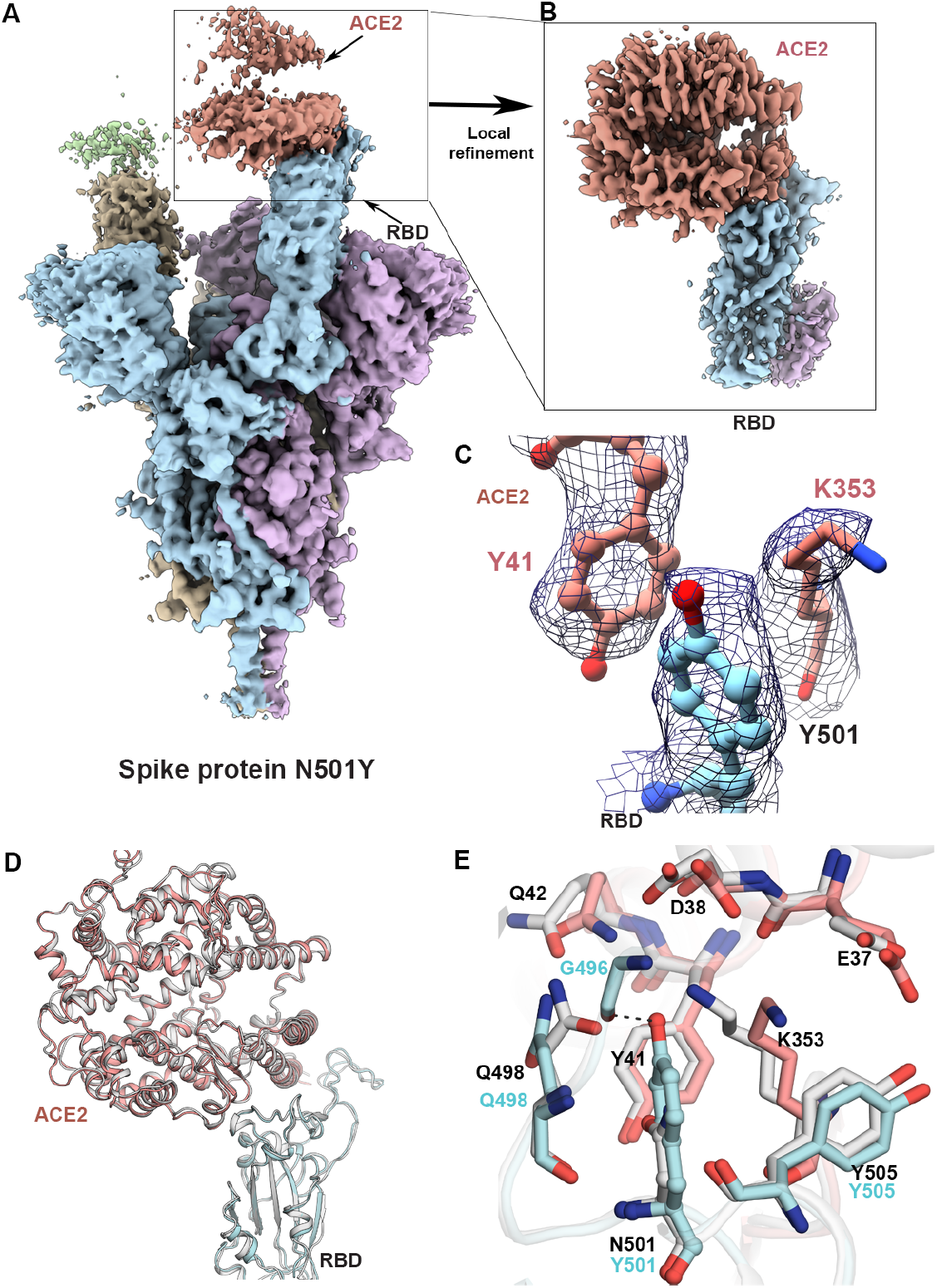
Structure of the SARS-CoV-2 N501Y mutant spike protein ectodomain bound to the ACE2 ectodomain. **(A)** Density map for the overall complex at the end of global structure refinement. The three spike protein protomers are colored in cyan, purple and yellow, with the density for the strongly and weakly bound ACE2 proteins in pale red and green, respectively. **(B)** Improved density map at the contact zone between the RBD and the strongly bound ACE2 protein ectodomain. **(C)** Visualization of density at the contact zone for Y501 in the RBD and residues Y41 and K353 in ACE2. **(D)**. Ribbon diagram with superposition of the unmutated and N501Y RBD-ACE2 complex (PDB ID: 7KMB). **(E)** Zoomed-in view of the interface, showing a superposition of the structures of unmutated and N501Y mutant spike proteins in complex with ACE2. The carbon atoms of residues in the N501Y mutant and ACE2 in our structure are colored in cyan and pale red respectively, while those in the structure of the complex between unmutated spike protein and ACE2 are in light gray.

### The N501Y mutation confers increased ACE2 binding affinity

Potent neutralization of SARS-CoV-2 has been achieved with a number of antibodies, including two recently reported examples V_H_ Fc ab8, and IgG ab1, both derived from a large human library of antibody sequences (*20, 21*). We compared the efficiencies of these two antibodies, as well as the ACE2 receptor ectodomain, to bind spike proteins with and without the N501Y mutation. We also determined the relative efficiency of neutralization of pseudoviruses expressing either the N501Y mutant or unmutated forms of the spike protein.

To test the influence of the N501Y mutation on ACE2 binding, we used a luciferase reporter to measure the infectivity of pseudotyped viruses presenting N501Y or unmutated spike proteins for cells overexpressing ACE2. The higher relative luminescence units (RLU) intensity from cells infected by the N501Y mutant (6000 ± 2000 RLU, mean ± standard deviation) compared to control viruses expressing the unmutated form (3000 ± 800 RLU) demonstrates that the N501Y mutation results in increased infectivity. To verify that this was a result of the increased binding strength of the mutant RBD to ACE2, we measured the binding parameters between ACE2 and of either unmutated or N501Y spike protein ectodomain trimers via biolayer interferometry (BLI). This revealed that the N501Y mutation confers an increase in affinity for ACE2, mainly driven by a reduction in dissociation rate constant (*k*_off_) (Figures 2A, B). We also measured the efficiency of exogenously added soluble ACE2-mFc proteins to neutralize unmutated and N501Y pseudoviruses (Figure 2C). The comparison of neutralization profiles show that the IC_50_ for neutralization of the N501Y mutant is lower. Taken together, these three results are consistent with the hypothesis that the greater infectivity of the N501Y mutant stems from improved binding to ACE2.

**Fig. 2.**
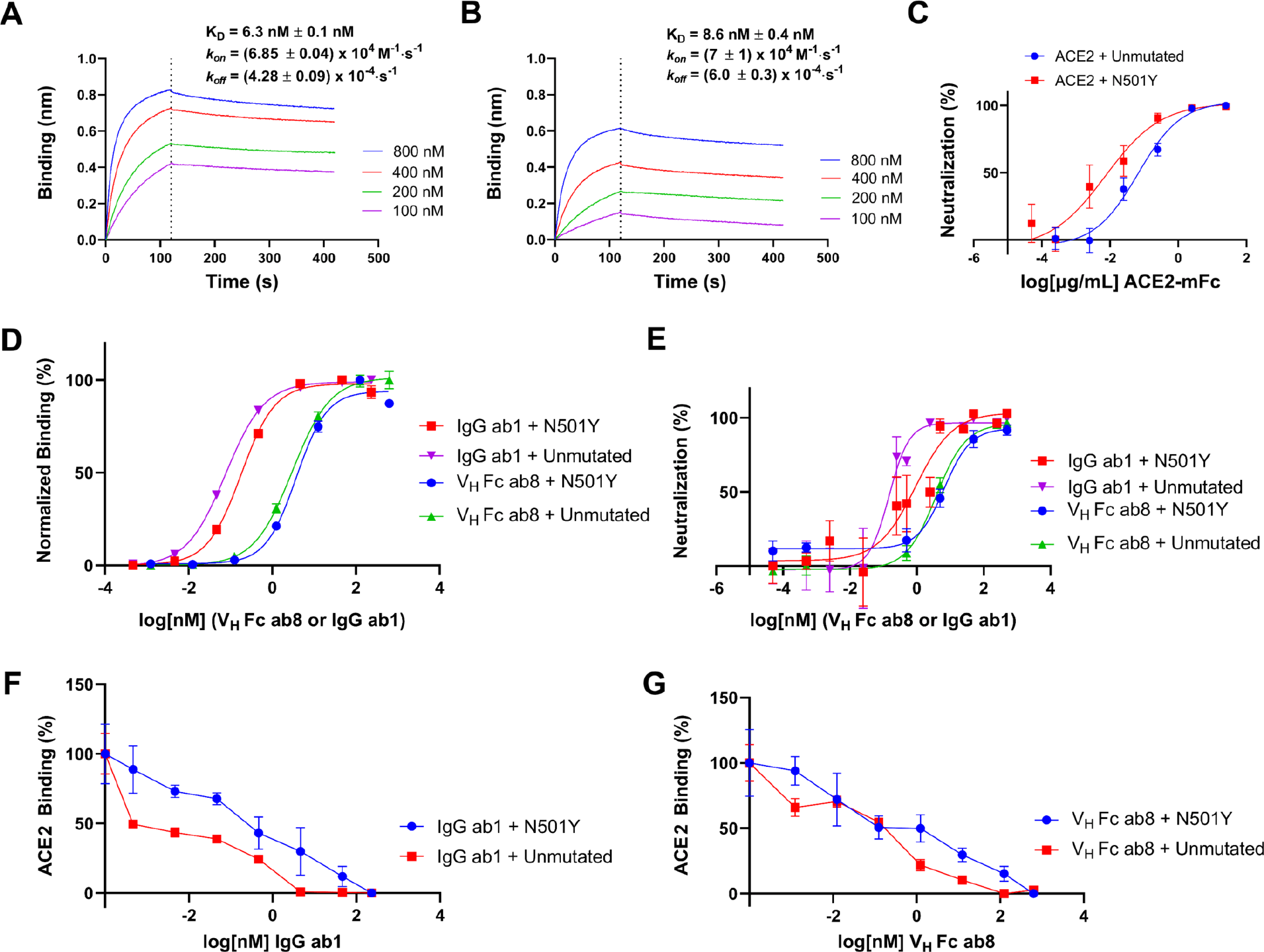
Analysis of ACE2, V_H_ Fc ab8 and IgG ab1 interactions with N501Y and unmutated spike. **(A,B)** Biolayer interferometry analysis of immobilized ACE2 binding by increasing concentrations of either N501Y (A) or unmutated (B) spike ectodomain. The extent of binding is shown as determined by the shift in wavelength (nm: nanometer). Biophysical parameters (K_D_, *k_on_, k_off_*) are shown as means ± standard deviations. **(C)** Analysis of N501Y or unmutated SARS-CoV-2 S-protein pseudo-typed virus neutralization by soluble ACE2-mFc. The IC_50_ of neutralization is greater for unmutated than for N501Y spikes as demonstrated by a one-tailed Welch test (P-value = 3E-5). **(D)** ELISA analysis of antibody interactions with either N501Y or unmutated spike ectodomain. **(E)** N501Y or unmutated SARS-CoV-2 S pseudo-typed virus neutralization by either V_H_ Fc ab8 or IgG ab1. **(F,G)** ELISA analysis of N501Y or unmutated SARS-CoV-2 spike ectodomain binding by soluble ACE2-mFc in the presence of serial dilutions of either (F) IgG ab1 or (G) V_H_ Fc ab8. ELISA experiments were done at least in duplicate while neutralization experiments were performed twice at least in duplicate, and the average values are shown. Error bars denote the standard error of the mean (SEM).

### N501Y has minimal effects on the binding and potency of two neutralizing antibodies with RBD epitopes

Next, we tested the effect of the N501Y mutation on the relative strengths of binding and neutralization potency of V_H_ Fc ab8 and IgG ab1 (Figures 2D-G). ELISA analysis of IgG ab1 and V_H_ Fc ab8 interactions with unmutated or N501Y spike ectodomains demonstrate the N501Y mutation has no significant effect on VH Fc ab8 binding but results in a slightly higher EC50 for IgG ab1 (Figure 2D). Second, competition experiments establish that IgG ab1 is more efficiently displaced by ACE2 when binding the N501Y mutant compared to unmutated ectodomains (Figure 2F), while the displacement of V_H_ Fc ab8 by ACE2 is similar for unmutated and N501Y mutant spike proteins (Figure 2G). This is further confirmed by negative stain experiments, where V_H_ ab8 interferes with ACE2 binding in both the unmutated and N501Y spikes (Figure S4). Consistent with these measurements, neutralization experiments carried out with V_H_ Fc ab8 shows that it can neutralize the N501Y mutant with a potency similar to that of the unmutated form, while IgG ab1 exhibits a slightly diminished neutralization potency for the N501Y mutant relative to pseudoviruses expressing the unmutated form (Figure 2E). Overall, binding and neutralization analyses show that the N501Y mutation results in enhanced ACE2 binding, minimal effects on the binding and potency of V_H_ Fc ab8, and a small reduction in the binding and potency of IgG ab1. To understand the effects of these antibodies at a structural level, we next determined cryo-EM structures of the complexes formed by V_H_ ab8 and Fab ab1 with the N501Y mutant spike protein ectodomain.

### Neutralizing antibodies bind N501Y spikes in different conformational states

Cryo-EM structural analysis of the complex formed between V_H_ ab8 and the N501Y spike protein ectodomain shows a single dominant conformation with two V_H_ ab8 fragments bound to RBDs in the down conformation and weak density for the other RBD, which is flexible and primarily in the up position (Figures 3A, S5). The global average resolution of the map is ~2.8 Å, with lower local resolution in the RBD regions, but local refinement yields maps of the V_H_ ab8– RBD interface at a resolution of ~3 Å (Figures 3B, S5). Cryo-EM density maps unambiguously show the location of residue 501 in the N501Y mutant spike protein ectodomains (Figure 3C). The interface between the RBD and V_H_ ab8 is well-defined with key interactions at the interface mediated by residues in the stretch between V483 and S494, along with a few other interactions contributed by non-contiguous RBD residues (Figures 3D, E). Residue 501 of the spike protein RBD is at the periphery of the footprint of ab8 and shows no evidence of interactions with the antibody. The presence of the mutation thus appears not to influence interactions between the RBD and V_H_ ab8.

**Fig. 3.**
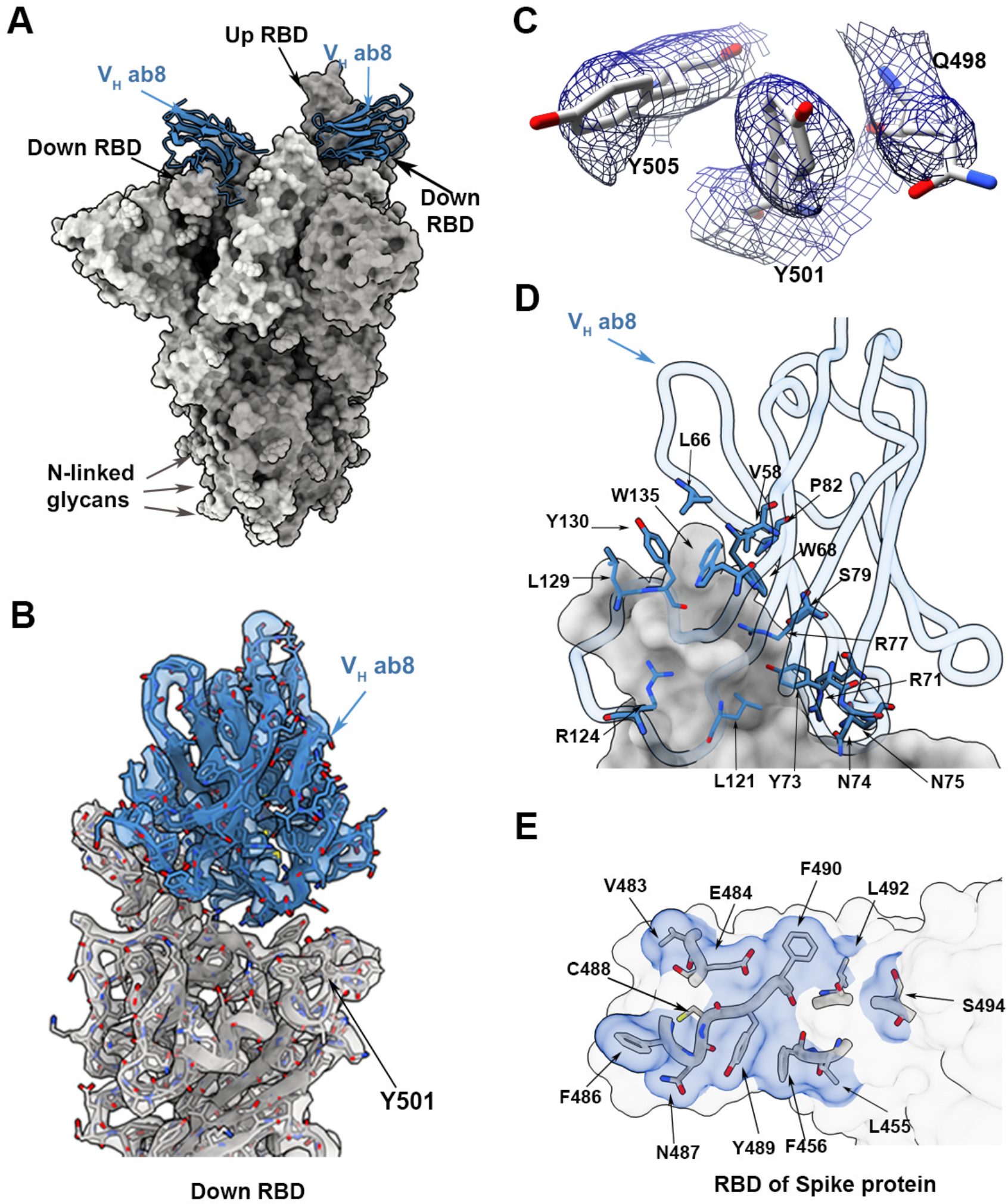
Structure of V_H_ ab8 bound to the N501Y mutant spike protein trimer. **(A)** Atomic model for the structure of the complex of V_H_ ab8 (blue) with the N501Y mutant spike protein ectodomain (grey). The structure has two RBDs in the down position with well-resolved densities for the bound V_H_ ab8. The third RBD is in the up position. **(B)** Cryo-EM density map after local refinement with fitted coordinates for the contact zone between the RBD and V_H_ ab8. **(C)** Density map in the region near 501 for the N501Y mutant spike protein ectodomain showing density for residues Q498, Y501 and Y505. **(D, E)** Close-up views of the contact zone between the RBD region and ACE2 highlighting residues involved.

Similar cryo-EM analyses of the complex between the mutated spike protein and Fab ab1 shows that in contrast to the V_H_ ab8 complex, Fab ab1 binding involves either two or three RBD domains, all being in the up position (Figures 4A, B, S6). Local refinement of the RBD–Fab ab1 interface improves the resolution to ~3 Å, enabling unambiguous placement of Y501 as well as the residues involved in the contact between the RBD and Fab ab1 (Figures 4C, D, S6). Residue 501 is at the periphery of the Fab ab1 footprint, with Ser 30 of Fab ab1 in a position to interact with the spike protein (Figures 4E, F). The N501Y mutation would thus be expected to have a small effect on the antibody binding epitope. Together, the cryo-EM structures are fully consistent with the studies presented in Figure 2 that show a small but significant effect of the N501Y mutation on Fab ab1 binding and neutralization, but with no measurable effects on V_H_ ab8 binding or neutralization.

**Fig. 4.**
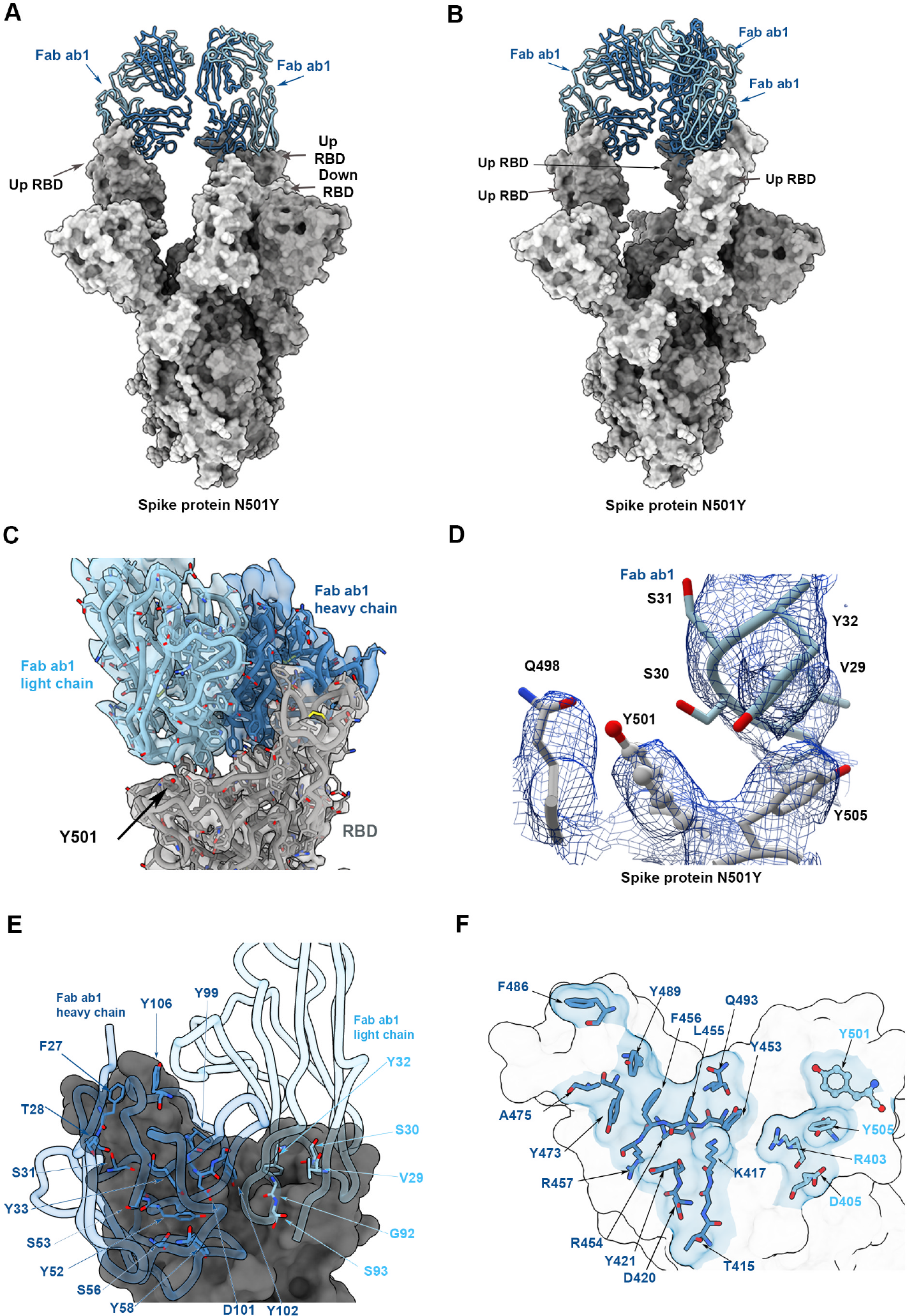
Structure of Fab ab1 bound to the N501Y mutant spike protein trimer. **(A, B)** Atomic models for the two predominant conformations of the spike protein (grey) observed with Fab ab1 (blue) bound to either two (A) or three (B) RBDs in the up position. **(C)** Cryo-EM density map after local refinement with fitted coordinates for the contact zone between the RBD and Fab ab1. **(D)** Density map in the region near 501 for the N501Y mutant spike protein ectodomain showing density for residues Q498, Y501 and Y505 in the spike protein and a loop in Fab ab1 that includes S30, the residue closest to Y501. **(E, F)** Close-up views of the contact zone between the RBD region and ACE2 highlighting residues involved.

## Discussion

Comparison of the structures reported here with those reported for the ACE2-RBD complex from earlier X-ray crystallography and cryo-EM studies enable visualization of the similarities and differences in the modes of binding (Figure 5). There are several regions such as the portion of the epitope in the vicinity of residue F486 that are shared across ACE2 and the two antibodies (Figures 5A-C). However, there are marked differences near residue 501, which is completely within the ACE2 footprint, at the very edge of the ab1 footprint, and well outside the ab8 footprint (Figures 5D-G). ACE2 binding has been observed only to RBDs in the up position, likely because of steric constraints in accommodating ACE2 in the down conformation. However, the stoichiometry of ACE2 binding to the trimeric spike can be variable. Negative stain experiments show that populations of spike proteins with one, two, or three ACE2 receptors bound are obtained (Figure S4) and consistent with the binding studies, we find that a higher number of ACE2 receptors bind N501Y spikes as compared to unmutated spikes when the incubation is carried out under similar conditions. In cryo-EM experiments, Fab ab1 also binds the RBD in only the up position (Figures 4A, B), but in contrast, the much smaller V_H_ ab8 fragment binds the RBD in both up and down positions (Figure 3A). Despite these differences, and the fact that ACE2, V_H_ ab8, and Fab ab1 each have distinctive directions of approach in their contact with the RBD, there is a good match in the RBD binding footprint between V_H_ ab8, Fab ab1, and ACE2 (Figures 5A-C), accounting for the potent neutralization by the V_H_ Fc ab8 and IgG1 ab1 antibodies (Figures 5D-G). (19). The location of residue 501 at the outer edge of the contact zone between RBD in the Fab ab1 complex and outside the zone of contact of V_H_ ab8 with RBD provides a structural rationale for the findings we describe here on the differential effects of the N501Y mutation on binding and neutralization by these two antibodies (Figure 5).

**Fig. 5.**
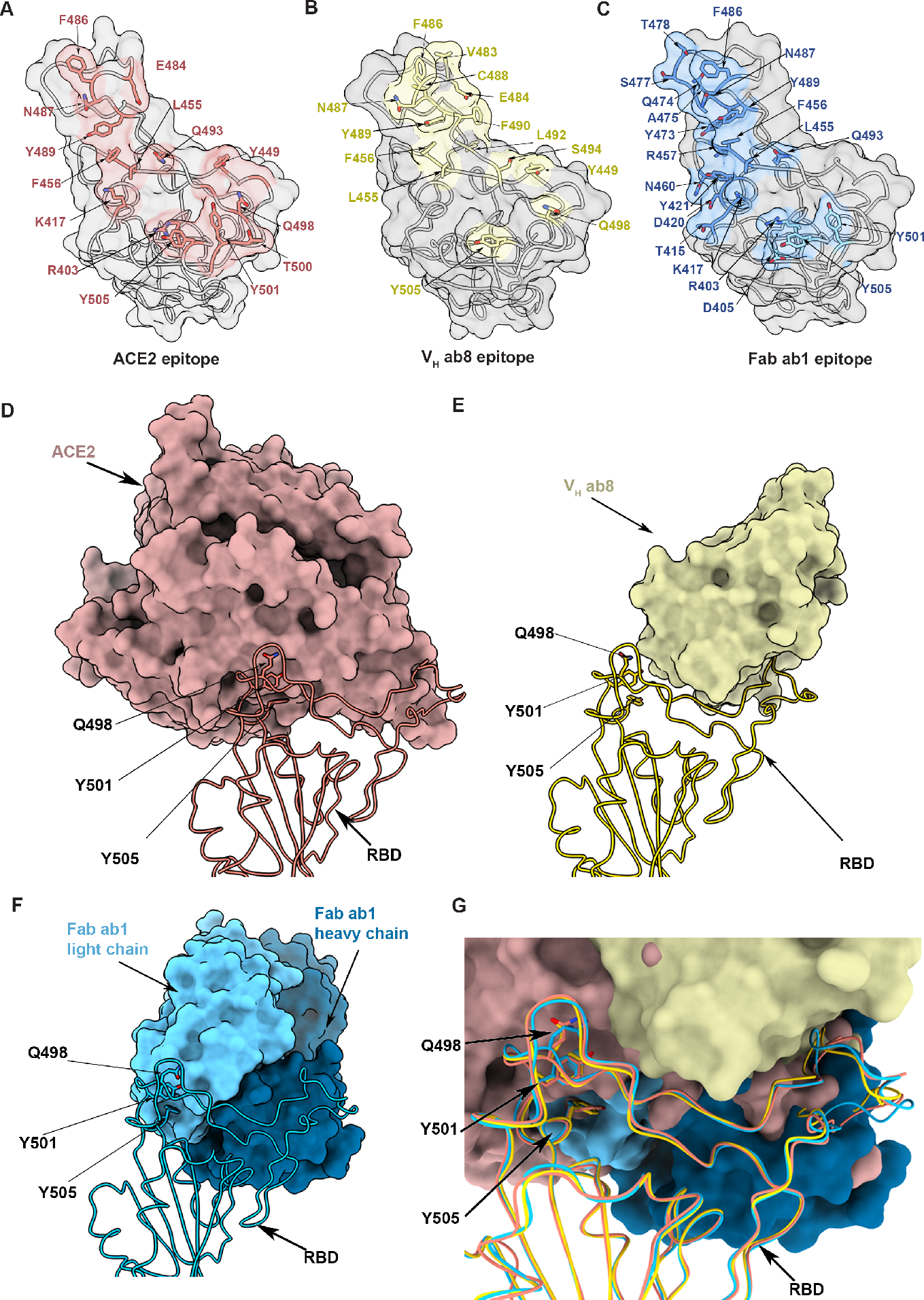
Comparison of the structures of complexes formed by the spike protein ectodomain with the ACE2 ectodomain, V_H_ ab8 and Fab ab1. **(A-C)** Open face views of the RBD from the vantage point of ACE2, V_H_ ab8 and Fab ab1 with the residues involved in contact shaded in red, yellow and blue, respectively. **(D-F)** Space-filling model view of ACE2 (D), V_H_ ab8 (E) and Fab ab1 (F) in contact with the RBD (structure shown in ribbon format). (G) Superposition of the structures of the complex of the RBD with ACE2, V_H_ Fc and Fab ab1 to show their relative footprints on the RBD surface.

Our studies with the N501Y mutant confirm the expectation that the rapid spread of the UK variant of SARS-CoV-2 is likely due to the viruses being more infectious. While there can be multiple origins for the increased infectivity, our biochemical and structural studies establish that the N501Y mutation results in increased ACE2 binding efficiency. Competition assays with a strongly neutralizing antibody show that it competes for binding with the spike trimer–ACE2 interaction in a concentration-dependent manner. Our results suggest that despite the higher infectivity of SARS-CoV-2 viruses carrying the N501Y mutation, the availability of the extended epitope surface on the RBD enables effective neutralization by V_H_ ab8 and Fab ab1. The footprints of these antibodies are comparable to that of other antibodies recently described (*22–25*), suggesting that at least some antibodies elicited by immunization with vaccines that are currently in production may also retain the ability to neutralize the N501Y mutant. With the continued spread of SARS-CoV-2, it appears likely that further mutations that enhance viral fitness will emerge. Cryo-EM methods to rapidly identify footprints of antibodies that are generated by current and future generations of vaccines could thus add a critical tool to the arsenal of efforts to prevent and treat COVID-19.

## Materials and Methods

### Cloning, Expression, and Purification of Recombinant Spike Protein Constructs

The wild type SARS-CoV-2 S HexaPro expression plasmid was a gift from Jason McLellan (7) and obtained from Addgene (plasmid #154754; http://n2t.net/addgene: 154754; RRID:Addgene_154754). The N501Y mutation was introduced by site-directed mutagenesis (Q5 Site-Directed Mutagenesis Kit, New England Biolabs). Successful subcloning and mutation were confirmed by Sanger sequencing (Genewiz, Inc.). Expi293F cells (ThermoFisher) were grown in suspension culture using Expi293 Expression Medium (ThermoFisher) at 37°C, 8% CO2. Cells were transiently transfected at a density of 3 × 10^6^ cells/mL using linear polyethylenimine (Polysciences). 24 h following transfection, media was supplemented with 2.2 mM valproic acid and expression carried out for 5 days at 37°C, 8% CO2. The supernatant was harvested by centrifugation and filtered through a 0.22-μm filter before loading it onto a 5 mL HisTrap excel column (Cytiva). The column was washed with 20 column volumes (CV) of wash buffer (20 mM Tris pH 8.0, 500 mM NaCl), followed by 5 CVs of wash buffer supplemented with 20 mM imidazole. The protein was eluted with elution buffer (20 mM Tris pH 8.0, 500 mM NaCl, 500 mM imidazole). Elution fractions containing the protein were pooled and concentrated (Amicon Ultra 100-kDa cut off, Millipore Sigma) for gel filtration (GF). Gel filtration was conducted using a Superose 6 10/300 GL column (Cytiva) pre-equilibrated with GF buffer (20 mM Tris pH 8.0, 150 mM NaCl). Peak fractions corresponding to soluble protein were pooled and concentrated to 4.5–5.5 mg/mL (Amicon Ultra 100-kDa cut off, Millipore Sigma). Protein purity was estimated as >95% by SDS-PAGE and protein concentration was measured spectrophotometrically (Implen Nanophotometer N60, Implen).

### Negative Stain Sample Preparation and Data Collection

For negative stain, purified S protein (0.05 mg/mL) was mixed with soluble ACE2 (0.05 mg/mL) and incubated on ice for 15 min. For the competition experiment, the S protein (0.05 mg/mL) was first incubated on ice with V_H_ ab8 (0.02 mg/mL) for 30 min, followed by addition of ACE2 (0.05 mg/mL) for another 30 min. Grids (Copper 200 or 300 mesh coated with continuous ultrathin carbon) were plasma cleaned using an H2/O2 gas mixture for 15 s in a Solarus II plasma cleaner (Gatan Inc.) or 10 s in a PELCO easiGlow glow discharge cleaning system (Ted Pella Inc.). The protein mixtures (4.8 μL) were applied to the grid and allowed to adsorb for 30 s before blotting away excess liquid, followed by a brief wash with MilliQ H2O. Grids were stained by three successive applications of 2% (w/v) uranyl formate (20 s, 20 s, 60 s). Negative stain grids were imaged using a 200 kV Glacios (ThermoFisher Scientific) transmission electron microscope (TEM) equipped with a Falcon3 camera operated in linear mode. Micrographs were collected using EPU at nominal 92,000x magnification (physical pixel size 1.6 Å) over a defocus range of −1.0 μm to −2.0 μm with a total accumulated dose of 40 e^−^/Å^2^.

### Cryo-EM Sample Preparation and Data Collection

For cryo-EM, both N501Y and unmutated SARS-CoV-2 spike ectodomain preparations were deposited on grids at a concentration of 2.25 mg/mL. Complexes were prepared by incubating spike ectodomain preparations with either ACE2 (residues 18–615, New England Biolabs), V_H_ ab8, or Fab ab1 at molar ratios of 1:1.25, 1:9, and 1:8 (spike trimer to binding partner), respectively. Incubations were performed for 20 min on ice prior to centrifugation at 14,000×g for 10 min. Grids were plasma cleaned using an H2/O2 gas mixture for 15 s in a Solarus II Plasma Cleaner (Gatan Inc.) before 1.8 μL of protein suspension was applied to the surface of the grid. Using a Vitrobot Mark IV (Thermo Fisher Scientific), the sample was applied to either Quantifoil Holey Carbon R1.2/1.3 Copper 300 mesh grids (N501Y spike alone and in complex with ACE2) or UltrAuFoil Holey Gold 300 mesh grids (N501Y spike in complex with V_H_ ab8 or Fab ab1) at a chamber temperature of 10°C with a relative humidity level of 100%, and then vitrified in liquid ethane after blotting for 12 s with a blot force of −10. All cryo-EM grids were screened using a 200 kV Glacios (ThermoFisher Scientific) TEM equipped with a Falcon4 direct electron detector followed by high-resolution data collection on a 300 kV Titan Krios G4 (ThermoFisher Scientific) TEM equipped with a Falcon4 direct electron detector in electron event registration (EER) mode. Movies were collected at 155,000x magnification (physical pixel size 0.5 Å) over a defocus range of −0.5 μm to −3 μm with a total dose of 40 e^−^/Å^2^ using EPU automated acquisition software (ThermoFisher).

### Image Processing

In general, all data processing was performed in cryoSPARC v.2.15 or v.3.0.1 (*26*) unless stated otherwise.

For negative stain data, motion correction and CTF estimation were performed in RELION v.3.1.1 (*27*). Particles were picked by crYOLO v.1.7.6 (*28*) with a general model (ftp://ftp.gwdg.de/pub/misc/sphire/crYOLO-GENERAL-MODELS/gmodel_phosnet_negstain_20190226.h5). After extraction, particles were imported into cryoSPARC and subjected to 2D classification and 3D heterogeneous classification. Final density maps were obtained by 3D homogeneous refinement.

For cryo-EM data, motion correction in patch mode (EER upsampling factor 1, EER number of fractions 40), CTF estimation in patch mode, reference-free particle picking, and particle extraction (extraction box size 640, Fourier crop to box size 320) were performed on-the-fly in cryoSPARC. After preprocessing, particles were subjected to 2D classification and 3D heterogeneous classification. The initial consensus maps were obtained by 3D homogeneous refinement. Then particles were re-extracted with box size 800 and then binned to 400. Final 3D refinement was done with per particle CTF estimation and aberration correction. Local refinements with a soft mask covering a single RBD and its bound V_H_ ab8 or ACE2 resulted in improvement of the binding interfaces. C3 symmetry expanded particles were used for local refinement of RBD and its bound Fab ab1. Overall resolution and locally refined resolutions were according to the gold-standard FSC (*29*).

### Model Building and Refinement

Coordinates of PDB 6WGJ and 7CH5 were used as initial models to build the V_H_ ab8 and Fab ab1, respectively. Individual domains of SARS-CoV-2 HexaPro S trimer (PDB code 6XKL) were docked into cryo-EM density using UCSF Chimera v.1.15 (*30*). Initial models were first refined against sharpened locally refined maps, followed by iterative rounds of refinement against consensus map in COOT v.0.9.3 (*31*) and Phenix v.1.19 (*32*). Glycans were added at N-linked glycosylation sites in COOT. Model validation was performed using MolProbity (*33*). Figures were prepared using UCSF Chimera, UCSF ChimeraX v.1.1.1 (*34*), and PyMOL (v.2.2 Schrodinger, LLC).

### Pseudovirus Neutralization Assay

SARS-CoV-2 S N501Y plasmid was obtained from SARS-CoV-2 S plasmid (HDM-IDTSpike-fixK) by site-directed mutagenesis (Q5 Site-Directed Mutagenesis Kit, New England Biolabs). SARS-CoV-2 S and SARS_CoV-2 S N501Y pseudotyped retroviral particles were produced in HEK293T cells as described previously (29). Briefly, a third-generation lentiviral packaging system was utilized in combination with plasmids encoding the full-length SARS-CoV-2 spike, along with a transfer plasmid encoding luciferase and GFP as a dual reporter gene. Pseudoviruses were harvested 60 h after transfection, filtered with 0.45 μm PES filters, and frozen. For cell-entry and neutralization assays, HEK293T-ACE2 cells were seeded in 96-well plates at 50,000 cells per well. The next day, pseudovirus preparations normalized for viral capsid p24 levels (Lenti-X™ GoStix™ Plus) were incubated with dilutions of the indicated antibodies, ACE2-mFc (SinoBiological), or media alone for 1 h at 37°C prior to addition to cells and incubation for 48 h. Cells were then lysed and luciferase activity assessed using the ONE-Glo™ EX Luciferase Assay System (Promega) according to the manufacturer’s specifications. Detection of relative luciferase units was carried out using a Varioskan Lux plate reader (ThermoFisher). Percent neutralization was calculated relative to signals obtained in the presence of virus alone for each experiment. The IC_50_ values were calculated using a four-parameter dose-response (sigmoidal) curve in GraphPad Prism (version 9 for Windows, GraphPad Software, San Diego, California USA, www.graphpad.com). This function provides the 95% confidence interval (95% CI) and standard error of the mean (SEM).

### Enzyme-Linked Immunosorbent Assay (ELISA)

100 μL of wild type or N501Y SARS-CoV-2 S protein preparations were coated onto 96-well MaxiSorp™ plates at 2 μg/mL in PBS overnight at 4°C. All washing steps were performed 5 times with PBS + 0.05% Tween 20 (PBS-T). After washing, wells were incubated with blocking buffer (PBS-T + 2% BSA) for 1 h at room temperature. After washing, wells were incubated with dilutions of V_H_ Fc ab8 or ACE2-mFc (SinoBiological) in PBS-T + 0.5% BSA buffer for 1 h at room temperature. After washing, wells were incubated with either goat anti-human IgG (Jackson ImmunoResearch) or goat anti-Mouse IgG Fc Secondary Antibody, HRP (Invitrogen) at a 1:8,000 dilution in PBS-T + 0.5% BSA buffer for 1 h at room temperature. After washing, the substrate solution (Pierce™ 1-Step™) was used for color development according to the manufacturer’s specifications. Optical density at 450 nm was read on a Varioskan Lux plate reader (Thermo Fisher Scientific). For ACE2 competition assays, experiments were conducted as described above with amendments. Serial dilutions of V_H_ Fc ab8 were incubated for 30 min at room temperature prior to the addition of 2.5 nM ACE2-mFc (SinoBiological). Wells were then further incubated for 45 min at room temperature. After washing, wells were incubated at a 1:8,000 dilution of Goat anti-Mouse IgG Fc Secondary Antibody, HRP (Invitrogen) in PBS-T + 0.5% BSA buffer for 1 h at room temperature. After washing, the substrate solution (Pierce™ 1-Step™) was used for color development according to the manufacturer’s specifications. Optical density at 450 nm was read on a Varioskan Lux plate reader (Thermo Fisher Scientific). For all experiments, controls for antibody–BSA interactions were performed. For competition assays, controls for anti-Mouse IgG Fc Secondary Antibody recognition of V_H_ Fc ab8 were performed. The EC50 values were calculated using a four-parameter dose-response (sigmoidal) curve in GraphPad Prism.

### Biolayer Interferometry (BLI)

The binding kinetics of SARS-CoV-2 trimers and human ACE2 was analyzed with the biolayer interferometer BLItz (ForteBio, Menlo Park, CA). Protein-A biosensors (ForteBio: 18– 5010) were coated with ACE2-mFc (40 μg/mL) for 2 min and incubated in DPBS (pH = 7.4) to establish baselines. Concentrations of 100 nM, 200 nM, and 400 nM spike trimers were used for association for 2 min followed by dissociation in DPBS for 5 min. The association (*k*_on_) and dissociation (*k*_off_) rates were derived from the sensorgrams fitting and used to calculate the binding equilibrium constant (KD).

## General

We thank Abel Gebresellasie and Brad Ross for their assistance with the maintenance of electron microscopes, and Peter Axerio-Cilies for helpful discussions. We thank TRIUMF and its personnel for help with the infrastructure supporting the operation of a Glacios electron microscope.

## Funding

This work was supported by awards to S.S. from a Canada Excellence Research Chair Award, the VGH Foundation, and Genome BC, Canada. W.L. and D.S.D. were supported by the University of Pittsburgh Medical Center. D.M. is supported by a CIHR Frederick Banting and Charles Best Canada Graduate Scholarship Master’s Award (CGS-M). J.-P.D. is supported by a Long-Term Fellowship (LT000988/2016-L) from the Human Frontier Science Program. J.W.S is supported by a CIHR Frederick Banting and Charles Best Canada Graduate Scholarships Doctoral Award (170893 – CGS-D) and a UBC President’s Academic Excellence Initiative Ph.D. Award (6817).

## Author contributions

This work was the result of a concerted team effort from all individuals listed as authors. D.M., K.L., S.S.S., S.Z. and J.W.S. collectively carried out all biochemical aspects including expression, production and purification of the spike proteins and antibody fragments. D.M. carried out the neutralization experiments. A.B., S.C., K.S.T. and J.-P.D. collectively carried out the experimental components of cryo-EM and electron microscopy including specimen preparation and data collection. X.Z. carried out all computational aspects of image processing and structure determination. X.Z., S.S.S. and S.S. interpreted and analyzed the cryo-EM structures. W.L. carried out the biolayer interferometry experiments. W.L. and D.S.D. provided the plasmids for V_H_ ab8 as part of a collaboration between the Subramaniam and Dimitrov laboratories on SARS-CoV-2. S.S. led and oversaw research on all aspects of the project and drafted the manuscript with input from all co-authors.

## Competing interests

All UBC authors declare no competing interests. Wei Li and Dimiter S. Dimitrov are coinventors of a patent, filed by the University of Pittsburgh, related to ab8.

## Data and materials availability

The density maps reported will be made publicly available in the Electron Microscopy Data Bank (EMDB: https://www.ebi.ac.uk/pdbe/emdb/) and atomic coordinates for the structures will be available at the Protein Data Bank (PDB: https://www.rcsb.org).

**Table 1.**
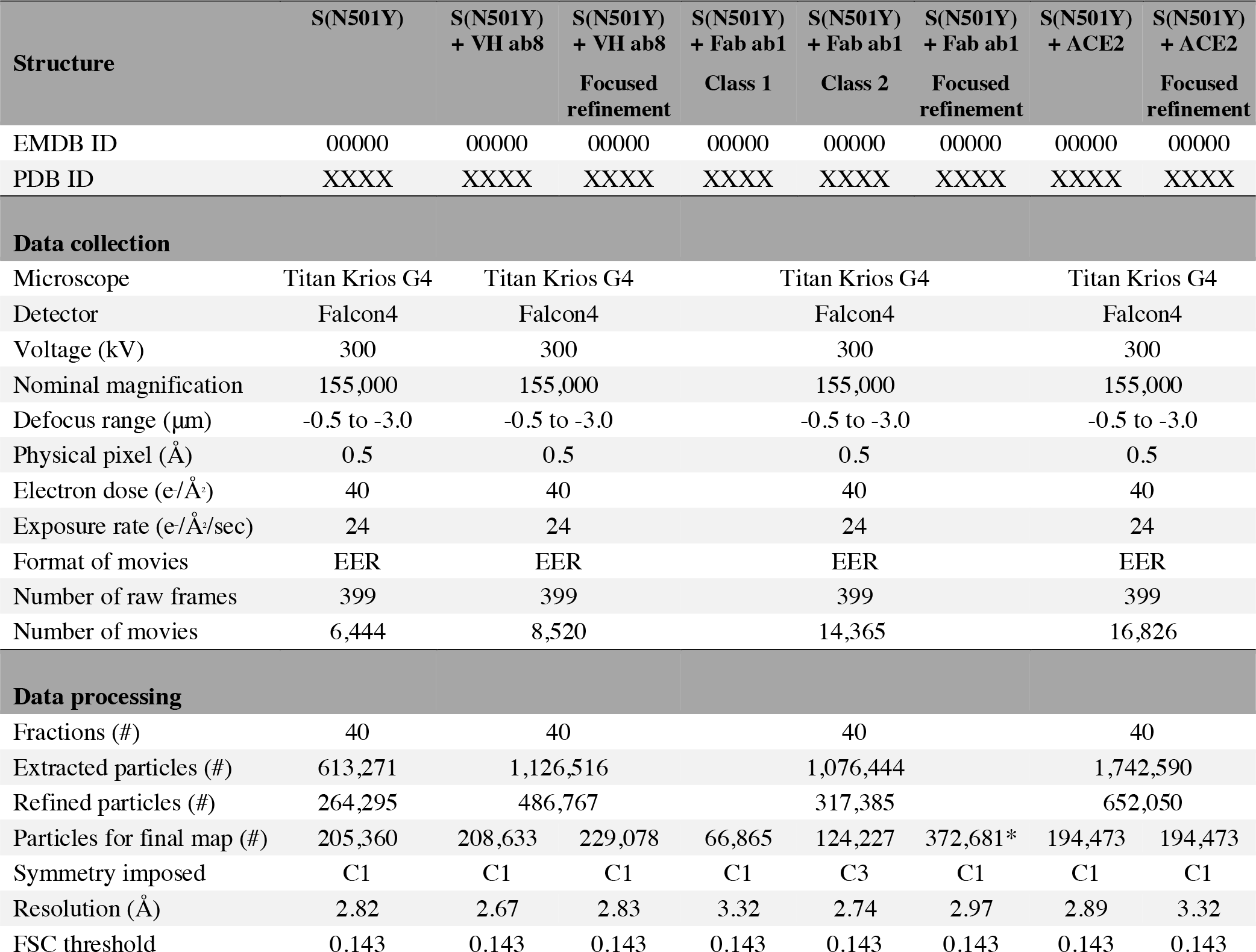
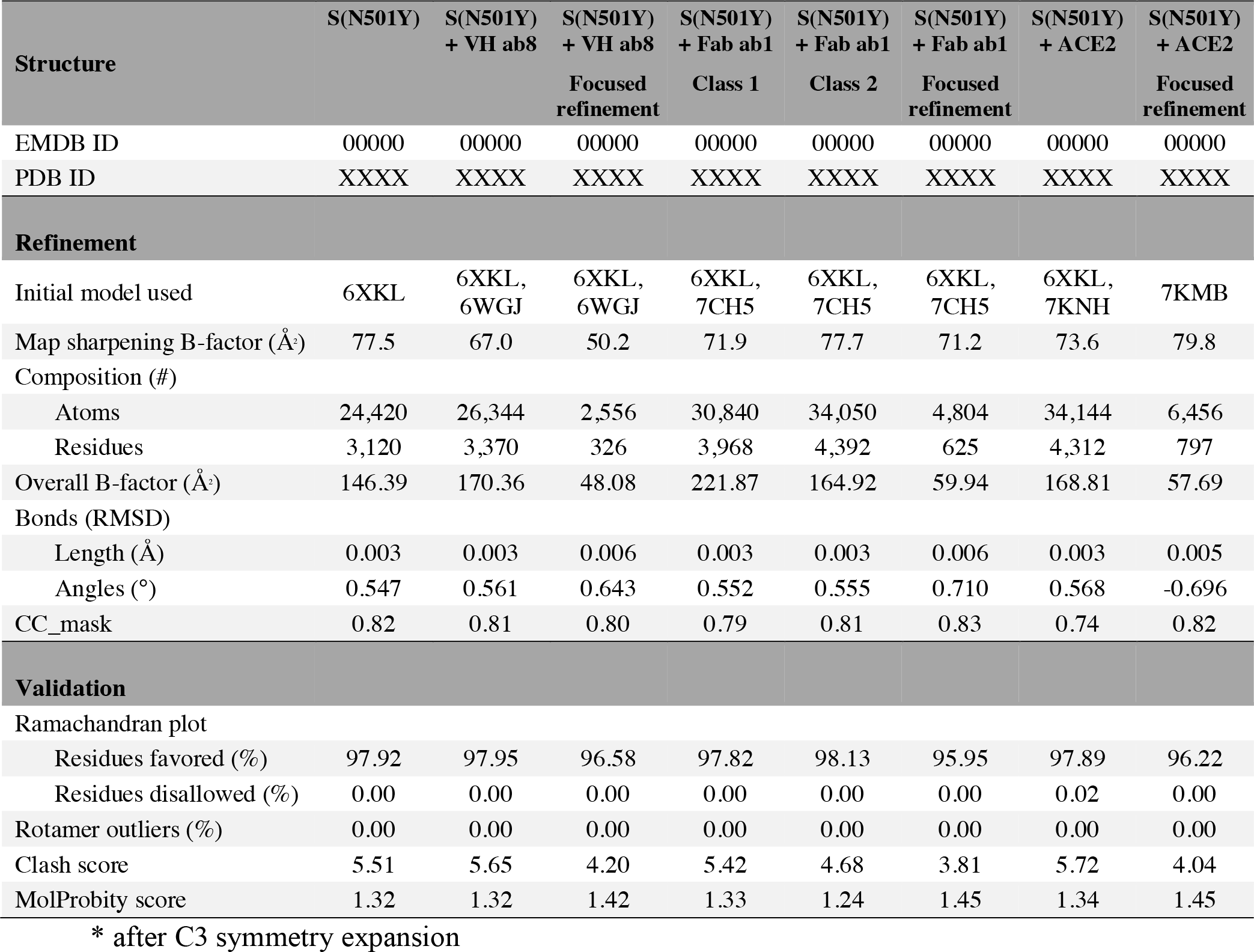
Data collection and processing parameters, refinement and validation statistics.

**Table 2.** Biophysical parameters.

| Assay | Biolayer interferometry |  |  | Pseudovirus Neutralization |  |  | ELISA |  |
| --- | --- | --- | --- | --- | --- | --- | --- | --- |
| | $k_{on}$ | $k_{off}$ | $K_D$ | $IC_{50}$<br>ACE2 | $IC_{50}$<br>IgG ab1 | $IC_{50}$<br>$V_H$ ab8 | $EC_{50}$<br>IgG ab1 | $EC_{50}$<br>$V_H$ ab8 |
| | $M^{-1} \cdot s^{-1}$ | $s^{-1}$ | nM | $\mu g/mL$ | nM | nM | nM | nM |
| Unmutated spikes | 7 | 6.0 | 8.6 | 0.066 | 0.14 | 3.76 | 0.0766 | 3.00 |
| ± Std. Dev.or CI 95% | ± 1 | ± 0.3 | ± 0.4 | 0.026–0.17 | 0.066–0.26 | 2.59–5.66 | 0.070–0.084 | 2.4–3.9 |
| N501Y spikes | 6.85 | 4.28 | 6.3 | 0.0074 | 0.79 | 6.61 | 0.181 | 3.75 |
| ± Std. Dev.or CI 95% | ± 0.04 | ± 0.09 | ± 0.1 | *–0.043 | 0.35–2.61 | 3.6–14.9 | 0.15–0.22 | 2.8–5.0 |

\* Lower bound not accurately determined

---

**Figure S1.**
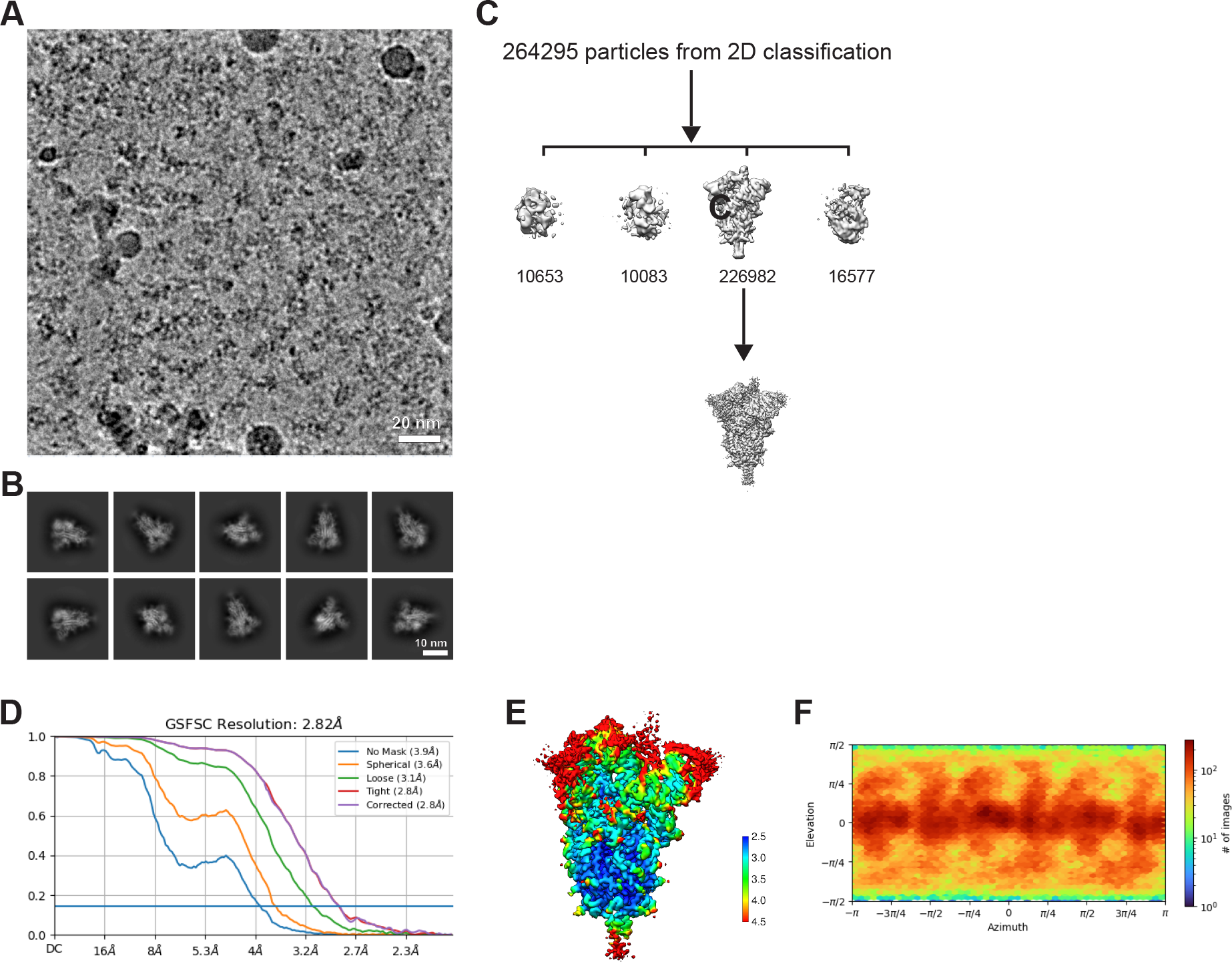
Cryo-EM data processing and validation for the N501Y spike protein ectodomains. (A) Representative micrograph. (B) 2D class averages. (C) 3D classification. (D) Gold-standard Fourier shell correlation (FSC) plot for global refinement. (E) Local resolution estimation. (F) viewing direction distribution.

**Figure S2.**
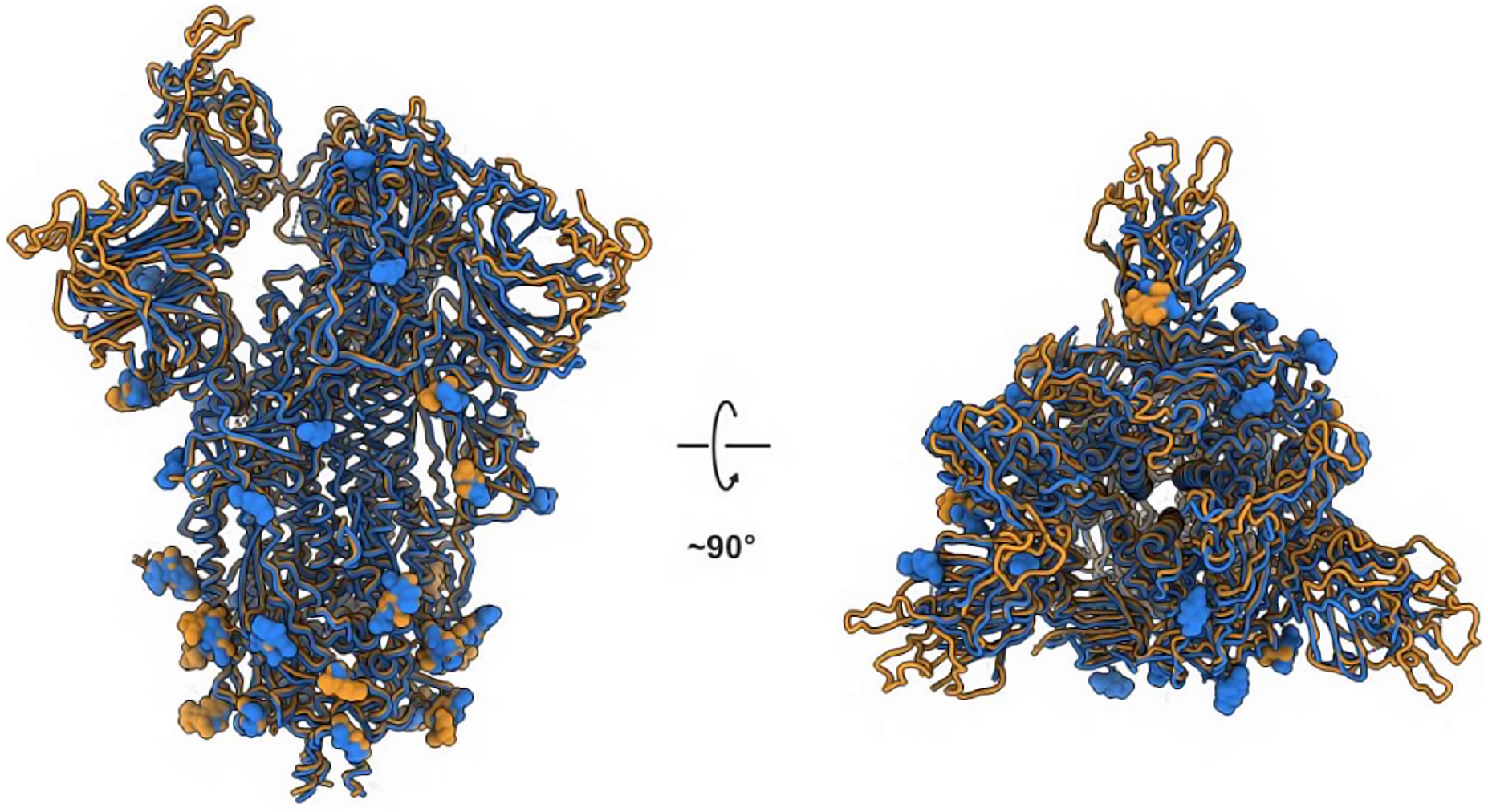
Superposition of the structure of the N501Y spike protein ectodomains (light orange) with the previously published structure of the unmutated construct (blue; PDB ID 6XKL).

**Figure S3.**
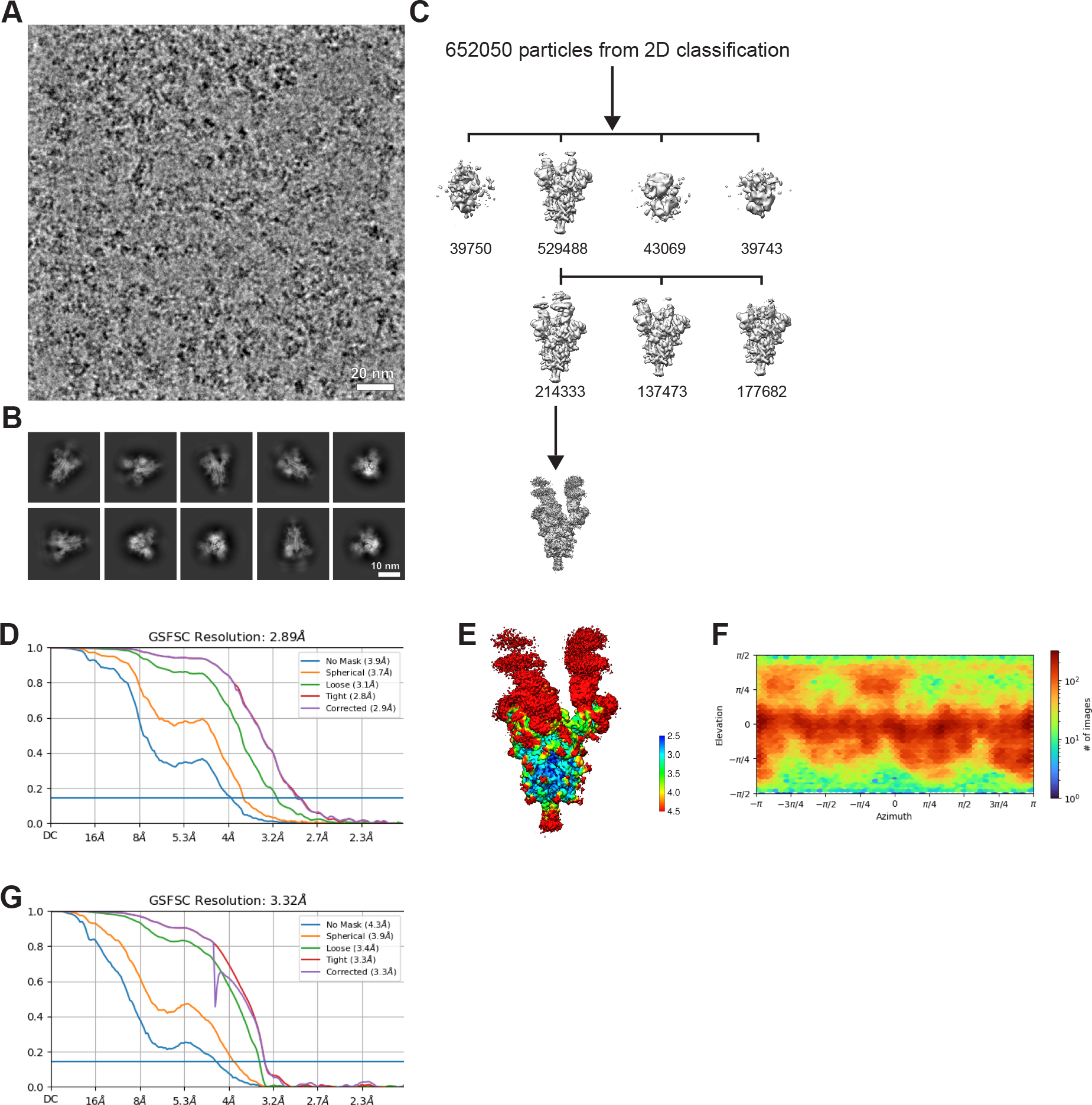
Cryo-EM data processing and validation for the complex between the N501Y spike protein ectodomain and the ACE2 ectodomain. (A) Representative micrograph. (B) 2D class averages. (C) 3D classification. (D) Gold-standard Fourier shell correlation (FSC) plot for global refinement. (E) Local resolution estimation. (F) viewing direction distribution. (G) FSC plot for local refinement.

**Figure S4.**
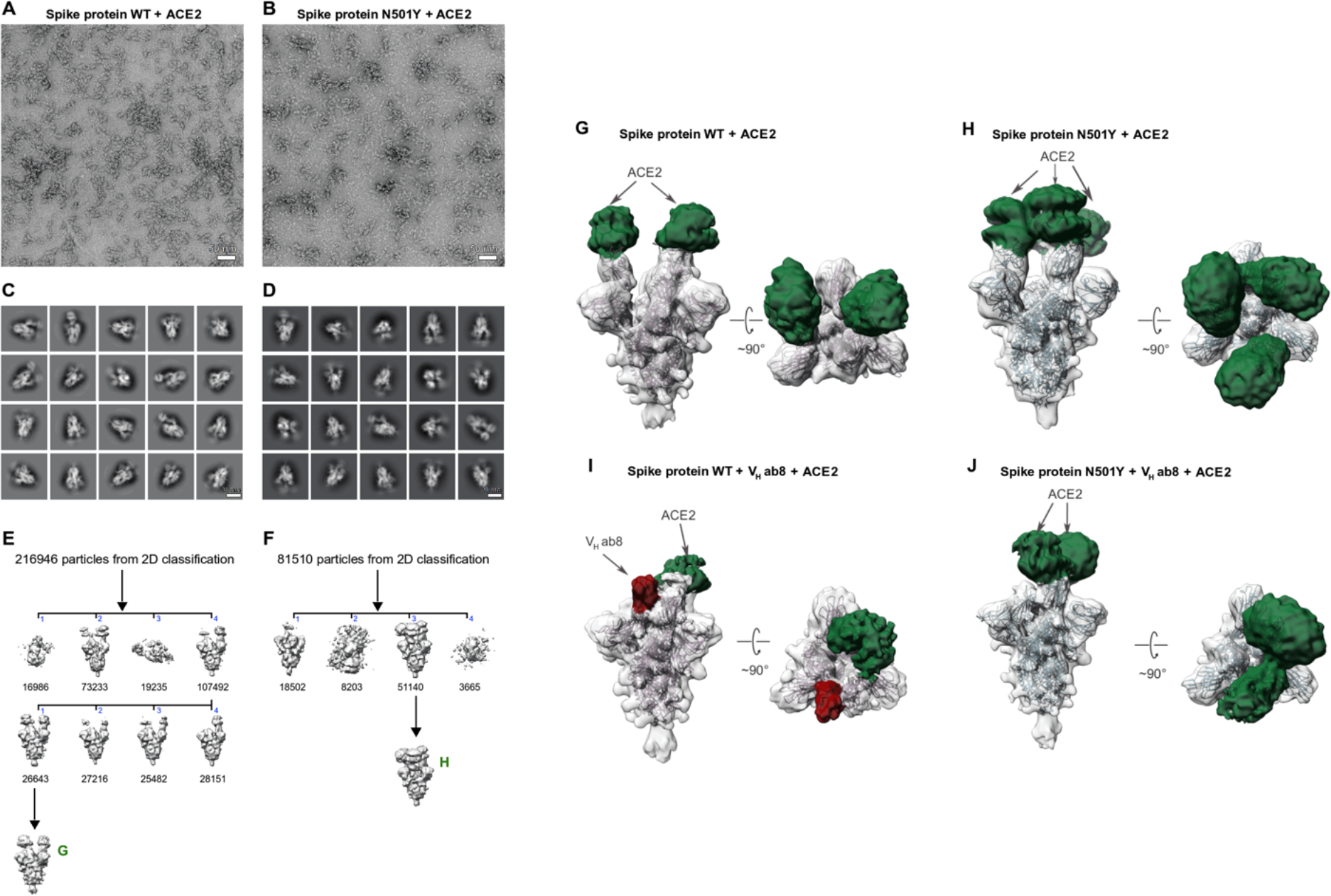
Negative stain EM reveals different ACE2 occupancies for unmutated and N501Y spikes (A, B) Representative micrograph selected from the total dataset for the unmutated (A) or N501Y (B) spike ectodomains in complex with ACE2. The concentrations of spike proteins and soluble ACE2 are the same for both unmutated and N501Y preparations. (C, D) 2D class averages corresponding to (C) the unmutated dataset (1355 images) and (D) N501Y dataset (1125 images), covering the same range of stain thickness. (E, F) Processing workflow. (E) For unmutated spikes, 3D classification reveals an occupancy of two RBD bound or less for the two most populated initial classes (50% and 34% of all particles from 2D classification). (F) For N501Y spikes, the most populated initial class (63%) has three RBD bound. (G, H) Final refinement of (G) unmutated spikes and (H) N501Y spikes. The density corresponding to bound soluble ACE2 is colored in green. The higher occupancy of ACE2 for N501Y spikes reflects a shift in the equilibrium stoichiometry, consistent with the higher affinity of N501Y spikes for ACE2. (I, J) Competition experiments. Spike ectodomains were first incubated with the V_H_ ab8 antibody fragment, then with soluble ACE2. The density corresponding to bound V_H_ ab8 is colored in red. The V_H_ ab8 antibody fragment competes with ACE2 binding as demonstrated by the reduced ACE2 occupancy in both (I) the unmutated spike (one RBD bound) and (J) the N501Y spike (two RBDs bound).

**Figure S5.**
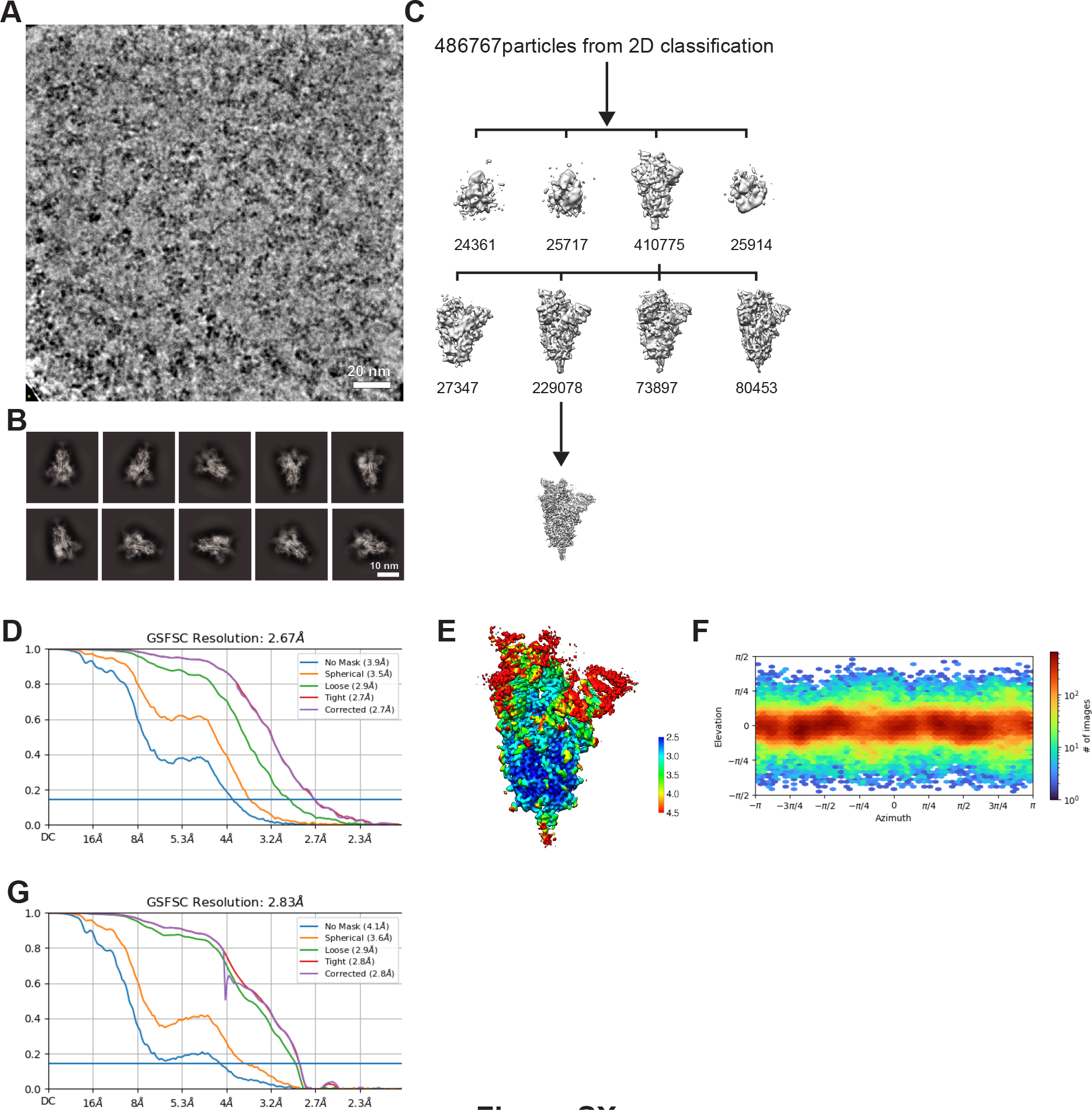
Cryo-EM data processing and validation for the complex between the N501Y spike protein ectodomain and V_H_ ab8. (A) Representative micrograph. (B) 2D class averages. (C) 3D classification. (D) Gold-standard Fourier shell correlation (FSC) plot for global refinement. (E) Local resolution estimation. (F) viewing direction distribution. (G) FSC plot for local refinement for the structure shown in Figure 3.

**Figure S6.**
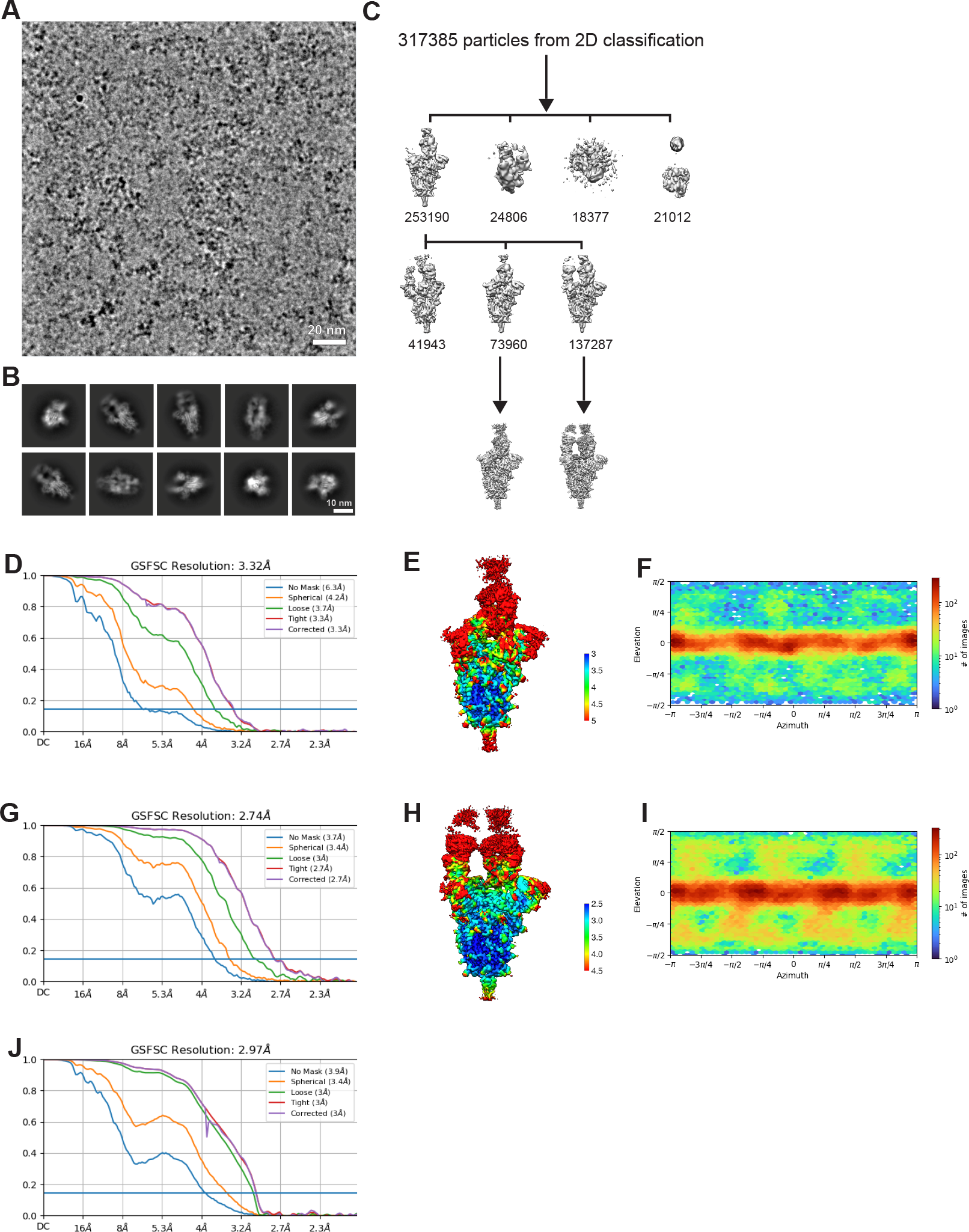
Cryo-EM data processing and validation for the complex between the N501Y spike protein ectodomain and Fab ab1. (A) Representative micrograph. (B) 2D class averages. (C) 3D classification resulting in two primary 3D classes. (D, G) Gold-standard Fourier shell correlation (FSC) plots for global refinement of the two classes. (E, F) Local resolution estimation corresponding to the maps shown in D and G, respectively. (F, I) viewing direction distribution corresponding to the maps shown in D and G, respectively. (J) FSC plot for local refinement of the structure shown in Figure 4.

## Notes

### Competing Interest Statement

All UBC authors declare no competing interests. Wei Li and Dimiter S. Dimitrov are co-inventors of a patent, filed by the University of Pittsburgh, related to ab8.

### Summary of Updates

The new results we have added to the revised manuscript include: 1)Cryo-EM structure of N501Y mutant spike protein in complex with ACE2 2)Cryo-EM structure of N501Y mutant spike protein in complex with Fab fragment of antibody ab1 3)Quantitative measurements of ACE2 binding affinity for unmutated and N501Y 4)Binding and neutralization studies with N501Y mutant proteins and viruses, respectively

